# Effects of live yeasts and their metabolic products on bumble bee microcolony development

**DOI:** 10.1101/2024.11.06.622134

**Authors:** Danielle Rukowski, Makena Weston, Rachel L. Vannette

## Abstract

Bumble bees can benefit from fungi, though the mechanisms underlying these benefits remain unknown and could include nutrition, resource supplementation, or pathogen protection. We tested how adding living yeasts or their metabolic products to *Bombus impatiens* diets in a factorial experiment affects microcolony performance, including survival, reproduction, and pathogen presence. We additionally assessed effects of yeast treatments on diet (nectar and pollen) chemical composition using untargeted metabolomics. Yeasts impacted microcolony reproduction and survival, but effects depended on source colony. Colonies containing the putative pathogen *Aspergillus* showed reduced reproduction, but yeast treatments reduced *Aspergillus* prevalence. Yeast treatments altered chemical composition of nectar and pollen, but most distinguishing compounds were unidentified. Our results suggest limited direct effects of yeasts via nutrition, resource supplementation, or modification of diets, instead suggesting that yeasts may benefit bees through interactions with the pathogens including *Aspergillus*. Overall, the effects of yeast supplementation are context-dependent, and more research is necessary to better understand the factors important in determining their impacts on bee hosts.

## Introduction

Insect-microbe associations are common and widely recognized for the role that microbes play in disease as well as the many beneficial functions they provide to their insect hosts. Most research has focused on the interactions between insects and bacteria, which are widespread and obligate in some insect systems (Feldhaar 2011, Kwong et al. 2017, Moran et al. 2005). However, microbial communities associated with insects are diverse, containing bacteria, fungi, viruses, and other taxa (Bessette & Williams 2022, Kaufman et al. 2000, Leigh et al. 2018, Scully et al. 2013). For some insect species, the importance of beneficial fungi has been recognized (Biedermann & Vega 2020), with fungi playing crucial roles in host biology, including acting as food, modifying host diet, or contributing compounds important to insect development. For example, several species of ants, termites, and beetles grow fungi in their nests as a food source (Mehdiabadi & Schultz 2009, Mueller & Gerardo 2002, Vanderpool et al. 2017). In addition, cactophilic *Drosophila* depend on fungi that inhabit the cactus wounds in which these flies breed and feed. These fungi detoxify toxic cactus compounds and produce sterols and vitamins essential to fly development (Starmer et al. 1986, Vega & Dowd 2005). Certain species of planthoppers, beetles, and bees also rely on associated fungi for the production of developmentally required sterols (Nasir & Noda 2003, Noda & Koizumi 2003, Paludo et al. 2018). Insect-associated fungi may also play a role in breakdown of complex food resources, including polysaccharides like cellulose, hemicellulose, or pectin (Li et al. 2012, Li et al. 2021, Vega & Dowd 2005).

Many bee species also host fungi, although the nature of these associations are just beginning to be understood (Rutkowski et al. 2023). A subset of these fungi are documented bee pathogens, including species in the genera *Aspergillus*, *Ascosphaera*, and *Vairimorpha*. However, these groups represent only a fraction of the fungal diversity associated with bee nests. Diverse groups of fungi are commonly isolated from the nests and guts of many bee groups, including honeybees, bumble bees, stingless bees, and solitary bees (Christensen et al. 2024, Dharampal et al. 2020, Echeverrigaray et al. 2021, Gilliam 1997, Rutkowski et al. 2023, Yun et al. 2018), many with unknown or beneficial impacts on bee health. Emerging evidence suggests that some bee species within the genus *Melipona* have lost bacterial symbionts and gained the yeast *Starmerella* within their gut instead (Cerqueira et al 2021) although the function of these yeasts remains unknown. For bumble bees (genus *Bombus*), yeasts have notable impacts on bee behavior and health. Foraging *Bombus* workers prefer to visit flowers inoculated with nectar yeasts over those without microbes (Schaeffer & Irwin 2014, Herrera et al. 2013, Yang et al. 2019), though these nectar yeasts were not found to impact colony development in *B. impatiens* (Schaeffer et al. 2017). Previous work in *Bombus impatiens* and *B. vosnesenskii* has found that adding nest-associated fungi to bee diets can improve bee survival and reproductive output, and help bees recover from negative effects of fungicide exposure (Rutkowski et al. 2022). Other work has found similar positive effects of adding flower or bee-associated fungi to the diets of *B. terrestris*, though these positive effects were dependent on the fungal species added (Pozo et al. 2020, Pozo et al. 2021).

The mechanisms behind these positive effects are unknown, but a few mechanisms are hypothesized (Steffan et al 2023). First, bees could derive nutrients from ingesting fungal cells, as has been implicated in the solitary bee genus *Osmia* (Dharampal et al. 2019, Steffan et al. 2019). Fungi may also produce vitamins and other metabolites important in bee diets or aid in the breakdown of dietary compounds. In the stingless bee *Scaptotrigona depilis*, a *Zygosaccharomyces* yeast provides sterols necessary for bee development, greatly improving larval survival (Paludo et al. 2018). In addition to nutritional impacts, fungi may directly suppress the growth of pathogens or improve bee immune function (Christensen et al. 2024, Pozo et al. 2020). A previous test of nutritional hypotheses using flower- and bee-associated fungal species found that different fungal species varied in their effects on *B. terrestris*, with some yeasts benefitting colony reproduction. However, the mode through which these fungi were provided had no impact (Pozo et al. 2020).

Here, we leverage a fungal community previously shown to benefit bumble bee fitness (Rutkowski et al. 2022) to test possible nutritional mechanisms through which fungi impact bee health. We hypothesize that fungi influence bumble bees through a combination of nutritional input from ingestion of fungal cells and fungal metabolites, and modification of nest provisions (stored nectar and pollen). To test this hypothesis, we factorially manipulated the presence of live fungal cells and fungal metabolites. We fed these solutions to microcolonies of the bumble bee *Bombus impatiens* and measured microcolony survival and reproductive success over the course of four weeks. If yeast effects are primarily chemically-mediated, the presence of live fungi or their metabolites (all yeast treatments) should benefit bumble bee survival and reproduction. If yeast effects are mediated via interactions between live yeasts and bees, only live yeasts (not metabolites alone) should benefit bees.

## Methods

### Bumble bee rearing conditions and experimental setup

We created 104 microcolonies of the commercially-reared bumble bee *Bombus impatiens* from 7 separate source colonies (Koppert, USA). Microcolonies are queenless colonies of bumble bee workers. In the absence of a queen, workers will lay eggs that develop into adult male offspring (Cnaani et al. 2002, Lopez-Vaamonde et al. 2007), allowing measures of microcolony survival as well as reproduction. *B. impatiens* is native to the eastern half of the United States, but commercial colonies are used throughout the US for pollination of greenhouse crops such as tomato and pepper (Velthuis et al. 2006). *Bombus impatiens* is often used in research as a model bumble bee species, and has been shown to benefit from addition of fungi in a previous study (Rutkowski et al. 2022). For this study, source colonies of *B. impatiens* were reared with nectar solutions provided by Koppert and sterilized honeybee-collected pollen (Koppert, USA). Pollen was sterilized using ethylene oxide to ensure that it was not contaminated with the bee pathogen *Ascosphaera apis*, which has previously been found in honeybee-collected pollen (Dharampal et al. 2020, Rutkowski et al. 2022) and can infect adult bumble bees (Maxfield-Taylor et al. 2015). Ethylene oxide gas was used due to the effectiveness of this sterilization method against fungi as well as its minimal side effects on bees compared to other common pollen sterilization methods (Strange et al. 2023). Although efforts were made to limit pathogen introduction via contaminated honeybee pollen, commercially reared colonies vary in the presence of pathogens and parasites (Strange et al. 2021).

Each microcolony consisted of five workers from a single source colony. Microcolonies were initially provided with a pollen ball, nectar, and wax pellets (Geelywax, China) to encourage reproduction. Pollen balls were created by mixing equal parts finely ground and sterilized honeybee-collected pollen (Koppert, USA) and sterile nectar. Nectar was created using a 2:2:1 ratio of glucose:fructose:sucrose, to a final total sugar concentration of 30%. Nectar was autoclaved prior to use to ensure sterility, after which non-essential amino acids (Cytiva, USA) were added to a final concentration of 5%. All microcolonies were initially fed with nectar inoculated with 7.5 ppm of the fungicide propiconazole (QualiPro, USA) for one week to reduce existing fungal abundance associated with each microcolony (Rutkowski et al. 2022).

Each microcolony was then assigned to a diet treatment consisting of 1) the presence or absence of live yeast cells and 2) the presence or absence of yeast metabolites. This created a total of four treatment groups (n=26 microcolonies/treatment). Treatment 1 included live yeast cells grown in artificial nectar for 5 days before being given to microcolonies (live yeasts + metabolites). Treatment 2 consisted of live yeast cells grown in artificial nectar for 5 days before cells were filtered out using a 0.2μm filter, leaving behind only modified nectar (metabolites only). Treatment 3 consisted of yeast cells newly suspended in artificial nectar (live yeasts only), and Treatment 4 consisted of sterile control nectar without yeast (no live yeasts, no metabolites). Yeasts *Debaryomyces hansenii, Starmerella sorbosivorans,* and *Zygosaccharomyces rouxii* were isolated from the honeypots of commercial *B. impatiens* colonies and were beneficial to bees in previous studies (Rutkowski et al 2022). These yeasts were grown separately and then added to the nectar of Treatments 1, 2, and 3 at a concentration of 1x10^4^ cells/mL for each species.

Microcolonies were kept on their treatments for four weeks, and nectar was replaced every three days (except in the case of Treatment 2, which was replaced every day to minimize yeast metabolism and modification of nectar). Pollen balls were made using these same treatment nectars and were replaced every three days until the start of reproduction, after which point colonies were given supplemental pollen as needed. Pollen balls were made once a week and stored at 4°C between replacements. Survival of workers in each microcolony was recorded daily, and time to emergence of the first adult offspring was tracked. To determine the impact of treatment on bumble bee feeding, nectar consumption over a three day period was measured for a subset of microcolonies (n=12/treatment) by weighing nectar cups before and after providing them to colonies. At the end of four weeks on their treatments, all surviving colonies were frozen and dissected to remove, count, and weigh all offspring (eggs, larvae, pupae, and adult males).

After microcolony creation, we noticed that microcolonies across all treatments that were created from source colonies that were older than one month (n=28) rarely produced offspring and died much sooner than other microcolonies. As many of these microcolonies died early into application of treatments, we did not include microcolonies sourced from old colonies in our analysis below. Treatments were equally represented in the dropped colonies, so that for all treatments, final sample size was equal (n=19 microcolonies/treatment).

### Pathogen presence and abundance

Because symptomatic fungal infection was observed in the larvae of some microcolonies, all microcolonies were screened for the presence and abundance of two common fungal pathogens (Strange et al. 2021). Diseased larvae from two microcolonies that exhibited fungal growth over their brood were plated on yeast media and resulting fungal growth was sampled and boiled in sterile water to extract DNA. DNA was amplified using PCR with the primers ITS86F (5’ GTGAATCATCGAATCTTTGAA 3’) and ITS4 (5’ TCCTCCGCTTATTGATATGC 3’) and the following run conditions: an initial denaturation at 95°C for 2mins, followed by 40 cycles of 95°C for 30s, 55°C for 30s and 72°C for 1min, and final extension step at 72°C for 10mins. Amplified DNA was sent to the UCDNA Sequencing Facility at UC Davis for Sanger sequencing using the same primers. The resulting sequences were identified using NCBI BLAST and both matched to *Aspergillus flavus* at 98.89% and 98.99% identity (Table S1). Therefore, all microcolonies were screened for pathogens using three primer sets targeting *Aspergillus flavus*, the *Aspergillus* genus, as well as the bee-specialist genus *Ascosphaera* (Evison & Jensen 2018), which has previously been detected in commercial *B. impatiens* colonies (Dharampal et al. 2020, Rutkowski et al. 2022).

To screen for pathogens, three workers in each microcolony were dissected to remove the midgut and hindgut. All three gut samples were pooled together and DNA was extracted using a PowerSoil Pro Kit (QIAGEN). To screen for the presence of *Aspergillus flavus*, we used the *A. flavus*-specific primers FLAVIQ1 (5’ GTCGTCCCCTCTCCGG 3’) and FLAVQ2 (5’ CTGGAAAAAGATTGATTTGCG 3’), using run conditions as in Sardiñas et al. 2011. To screen for the presence of the genus *Aspergillus,* we performed PCR using the genus-specific primers ASAP1 (5’ CAGCGAGTACATCACCTTGG 3’) and ASAP2 (5’ CCATTGTTGAAAGTTTTAACTGATT 3’), using run conditions as in Sugita et al. 2004. To screen for *Ascosphaera*, the genus-specific primers AscoALL-1 (5’ GCACTCCCACCCTTGTCTA 3’) and AscoALL-2 (5’ GAWCACGACGCCGTCACT 3’) were used, using run conditions as in James & Skinner 2005.

Multiple colonies tested positive for the presence of *A. flavus*, and abundance was quantified using qPCR on samples using the same primers and qPCR cycle conditions specified in Sardiñas et al. 2011. We used a custom plasmid (Eurofins Genomics Blue Heron) containing a known concentration of *A. flavus* amplicon DNA to convert from Cq values to total abundance (Table S2).

### Chemical analysis of yeast modifications to diet

To understand effects of yeast growth on the chemical composition of diet, nectar and pollen were collected from each microcolony three days after treatment application and a subset of these samples (n=5 per treatment, from two source colonies) were submitted to the West Coast Metabolomics Center at UC Davis for untargeted metabolomics using GC-TOF MS. To prepare nectar samples, 30uL of nectar was extracted using 1mL of a 5:2:2 methanol:chloroform:water solution. Samples were then vortexed for 10s, shaken for 5 mins, and centrifuged for 2 mins at 14,000rcf. Each 50uL aliquot of the resulting supernatants was dried. For pollen samples, 4.0±0.3mg of pollen was extracted using 1mL of the same solution as for nectar. Samples were then sonicated for 1hr with 1.6mm steel balls, shaken for 5mins, and centrifuged for 2 mins at 14,000rcf. From the resulting supernatant, 450uL aliquots were dried.

Aliquots of 0.5uL of prepared samples, resuspended in MSTFA, were injected into a 7890A GC coupled with a LECO Pegasus IV TOF MS, onto a RESTEK RTX-5SIL MS column with an Intergra-Guard at 275°C with a helium flow of 1 mL/min using a splitless method. Samples were held in the GC at 50°C for 1 min before heating to 330°C at a rate of 20°C/min, followed by sample holding for an additional 5 mins. The transfer line was heated to 280°C and the EI ion source to 250°C. The MS collected data from 85m/z to 500m/z at an acquisition rate of 17 spectra/s. Data preprocessing was performed using ChromaTOF version 2.32, with automatic mass spectral deconvolution and peak detection using a 5:1 signal:noise ratio. Preprocessed chromatograms were then identified using the BinBase database, using the associated BinBase algorithm. Peak height was used to report compound quantification and used in statistical analysis (below).

### Statistical analysis

All data analysis was carried out in R version 4.2 (R Core Team 2022). We tested if the presence of live yeasts (presence/absence), metabolites (presence/absence), or source colony influenced microcolony parameters. We tested for differences in the likelihood that a microcolony produced any offspring across treatments using a generalized linear model with a binomial distribution. Offspring abundance (total and for each life stage) between treatments was analyzed using a generalized linear model with a negative binomial distribution (‘MASS’ package, Venables & Ripley 2022). The effect of worker survival on total offspring abundance was analyzed using a generalized linear model with a negative binomial distribution. This survival term was calculated as the average number of workers alive during the entire duration of the microcolony life span. We compared the average mass of offspring (separately for each life stage) across treatments using ANOVA. Larval mass was not normally distributed and was square-root transformed prior to analysis. As few microcolonies produced pupae and adult males, the three way interaction between treatments and source colony was not included in these analyses. Differences in offspring composition (proportion of developmental stages) between treatments was analyzed using a chi-square test, run on a contingency table of proportions of each offspring stage (eggs, larvae, pupae, and adults) by treatment.

Survival of microcolony workers was analyzed using a Cox proportional hazards model for recurrent events (‘survival’ package, Therneau 2022), with live yeast presence, metabolite presence, and source colony as predictors and microcolony ID as a random effect. Per-bee nectar consumption was compared between treatments using ANOVA.

To test if the presence and abundance of presumed pathogens differed between yeast treatments, *Aspergillus* or *A. flavus* presence across treatments and source colonies was analyzed using a generalized linear model with a binomial distribution, while abundance of *A. flavus* was analyzed using an ANOVA. For these models, live yeast presence, metabolite presence, source colony, and all interactions between factors were included as predictors. Prior to all subsequent analyses of *Aspergillus* impacts on offspring production, data was subset to only include source colonies that tested positive for *Aspergillus*. The impact of *Aspergillus* presence on offspring presence was analyzed using a generalized linear model with a binomial distribution, with *Aspergillus* presence in a microcolony, live yeast presence, metabolite presence, and their interaction as predictors. The impact of *Aspergillus* on offspring abundance was analyzed using a generalized linear model with a negative binomial distribution, with *Aspergillus* presence in a microcolony, live yeast presence, metabolite presence, and their interaction as predictors. The impact of *Aspergillus* on survival was analyzed using a Cox-proportional hazards model, with live yeast presence, metabolite presence, *Aspergillus* presence within a microcolony, and their interactions as predictors, and microcolony ID as a random effect. Only *Aspergillus* impacts on offspring production and worker survival were analyzed, as *A. flavus* was present in appreciable amounts from only five microcolonies, and *Ascosphaera* was not detected in any microcolonies.

Metabolomics data was filtered and normalized using MetaboAnalyst 6.0 (Pang et al. 2021). Nectar and pollen samples were processed and analyzed separately, with live yeast presence and metabolite presence as main and interactive factors. According to standard MetaboAnalyst filtering recommendations, features were first filtered using median to remove low-value features, and then using interquartile range to remove features that remained constant throughout the dataset. Raw data contained 564 features, and after filtering contained 423 features. Data were normalized by median and auto scaled (mean-centered and divided by the standard deviation of each variable) before analysis to achieve a more normal distribution. The ANOVA2 function in MetaboAnalyst was used to test for differently abundant compounds between treatment groups, with the false discovery rate used for multiple testing p-value correction. For subsequent analyses, normalized data was imported into R. We ran a redundancy analysis (‘vegan’ package, Oksanen et al. 2022) to test for differences in metabolite profile between treatment groups, followed by permutation tests to determine significance (999 permutations). Random forest analysis (‘randomForest’ package, Liaw & Wiener 2002) was run to determine accuracy in classifying samples to treatments based on metabolomic profiles, using 10,001 trees per model. Results were visualized using the packages ‘ggplot2’ (Wickham 2016) and ‘viridis’ (Garnier et al. 2023).

## Results

### Yeast effects on microcolony reproduction vary with source colony

80% of microcolonies produced at least one offspring, with a mean of 14.4 offspring per microcolony (range 0-60 offspring). The probability that a microcolony produced at least one offspring was impacted by source colony (*Χ* ^2^ = 12.9, df = 6, p = 0.045), but was unaffected by metabolite presence (*X* ^2^ = 0.16, df = 1, p = 0.69), live yeast presence (*X* ^2^ = 0.59, df = 1, p = 0.44), or any interactions between factors (p > 0.05 for all comparisons). Microcolonies that survived longer were more likely to produce offspring (*X* ^2^ = 7.1, df = 1, p = 0.008).

Total offspring abundance (eggs, larvae, pupae, and adult males) was affected by both live yeast presence and metabolite presence, but these effects were dependent on source colony (live yeast presence * source colony interaction: *X* ^2^ = 26.8, df = 6, p < 0.001, metabolite presence * source colony interaction: *X* ^2^ = 19.2, df = 6, p = 0.004, Figure 1). For four of the seven source colonies, fungal treatment in some form was beneficial to offspring production, while for the other three colonies, treatments were either detrimental or had no effect. Offspring abundance was also higher in microcolonies that lived longer (*X* ^2^ = 22, df = 1, p < 0.001).

**Figure 1.**
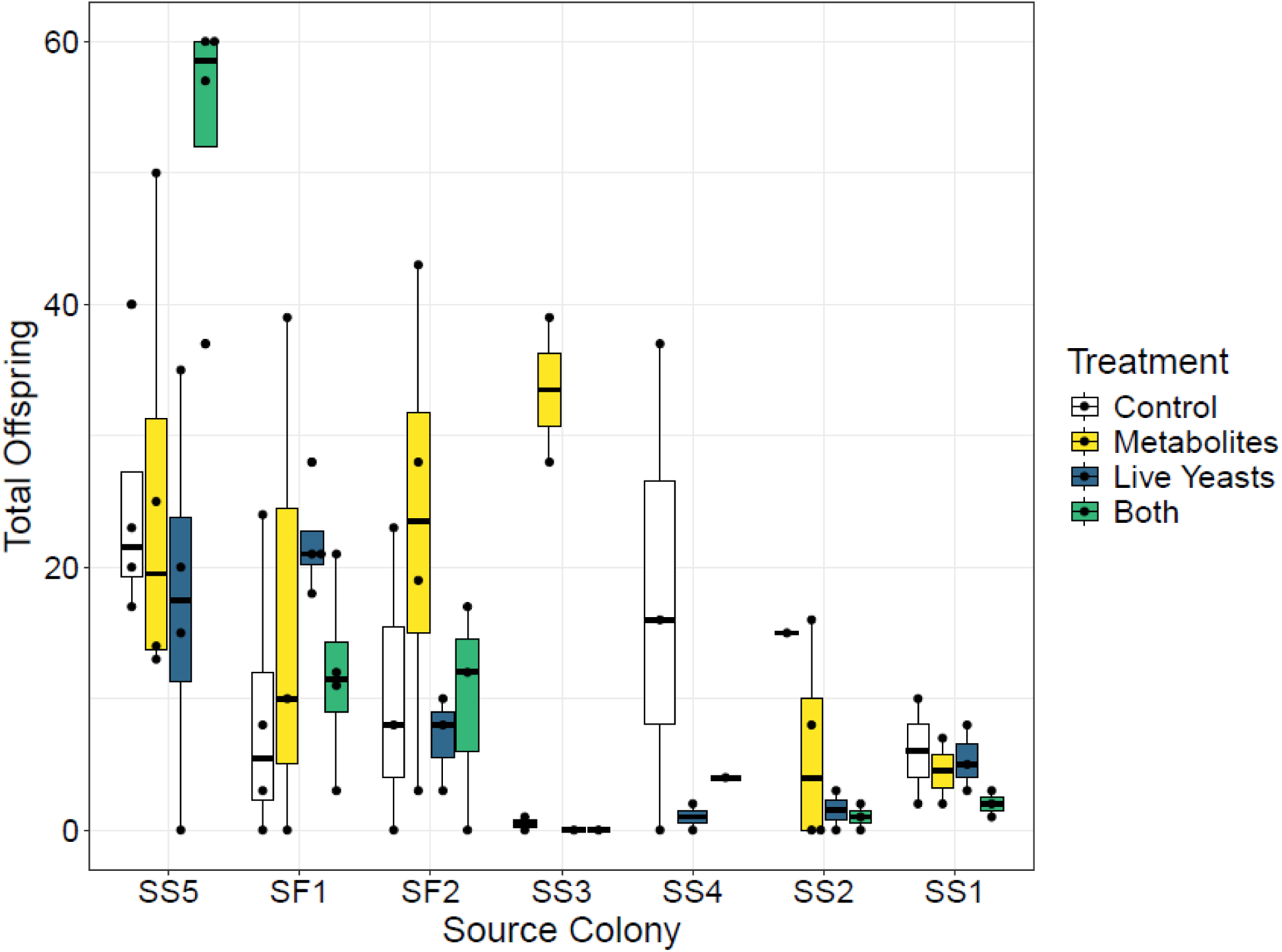
Total number of offspring (eggs, larvae, pupae, and adult males) produced by *B. impatiens* microcolonies exposed to different fungal treatments, faceted by source colony. Treatment effects were dependent on the source colony the microcolony was from (live yeast presence * source colony interaction: *X* ^2^ = 26.8, df = 6, p < 0.001, metabolite presence * source colony interaction: *X* ^2^ = 19.2, df = 6, p = 0.004).

Considering each life stage separately, the abundance of eggs, larvae, pupae, and adult males was also dependent on the interactions between treatments and source colonies (Table 1). The mass of most life stages was unaffected by treatments or source colony, except for larval mass, which was lower in microcolonies given yeast metabolites (Table S3). Offspring composition was significantly different between treatments (*X* ^2^ = 19.2, df = 9, p = 0.02, Figure 2), with microcolonies receiving only live yeast cells containing compositionally more larvae and males than other treatments.

**Figure 2.**
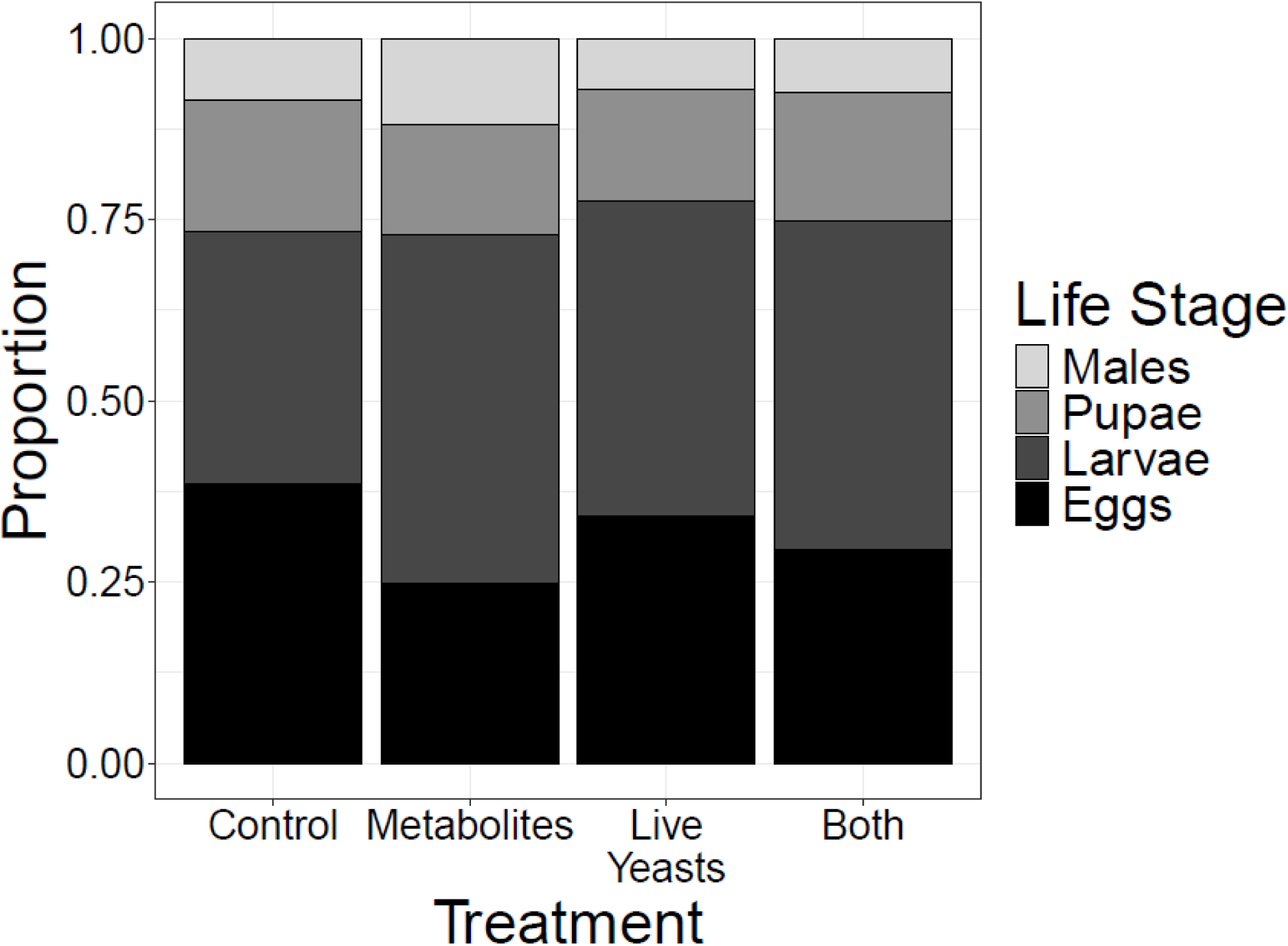
The life-stage makeup of the offspring produced by *B. impatiens* microcolonies exposed to different fungal treatments. Offspring composition was different between treatments, with the live yeasts only treatment having a higher proportion of larvae and males (*X* ^2^ = 19.2, df = 9, p = 0.02).

**Table 1.** Effects of yeast treatments and source colony on microcolony offspring abundance, separated by life stage.

| Response variable | Predictor variable | $X^2$ value | p-value |
| --- | --- | --- | --- |
| Egg number | Live yeast presence | 2.96 | 0.086 |
|  | Metabolite presence | 0.01 | 0.92 |
|  | **Source colony** | **47.7** | **< 0.001** |
|  | Live yeast presence * Metabolite presence | 0.01 | 0.92 |
|  | **Live yeast presence \* Source colony** | **17.1** | **0.009** |
|  | **Metabolite presence \* Source colony** | **18.7** | **0.005** |
|  | Live yeast presence * Metabolite presence * Source colony | 9.62 | 0.087 |
| Larvae number | Live yeast presence | 0.53 | 0.47 |
|  | Metabolite presence | 1.64 | 0.20 |
|  | **Source colony** | **61.5** | **< 0.001** |
|  | Live yeast presence * Metabolite presence | 0.39 | 0.53 |
|  | **Live yeast presence \* Source colony** | **17.2** | **0.009** |
|  | Metabolite presence * Source colony | 11.5 | 0.075 |
|  | Live yeast presence * Metabolite presence * Source colony | 2.32 | 0.80 |
| Pupae number | Live yeast presence | 2.00 | 0.16 |
|  | Metabolite presence | 2.29 | 0.13 |
|  | **Source colony** | **24.2** | **< 0.001** |
|  | Live yeast presence * Metabolite presence | 0.12 | 0.73 |
|  | **Live yeast presence \* Source colony** | **12.7** | **0.049** |
|  | Metabolite presence * Source colony | 12.4 | 0.054 |
| Male number | Live yeast presence | 0.0002 | 0.99 |
|  | Metabolite presence | 0.97 | 0.33 |
|  | **Source colony** | **23.5** | **< 0.001** |
|  | Live yeast presence * Metabolite presence | 3.65 | 0.056 |
|  | **Live yeast presence \* Source colony** | **15.7** | **0.015** |
|  | **Metabolite presence \* Source colony** | **16.8** | **0.01** |

### Effects of treatments on worker survival varies by source colony

Worker survival was affected by the three-way interaction between live yeast presence, metabolite presence, and source colony (*X* ^2^ = 25.8, df = 5, p < 0.001, Table 2). Workers in microcolonies from four source colonies survived longer when given fungal treatments in some form, though the most beneficial mode of application varied across source colonies. For the other three source colonies, fungal treatments had either neutral or negative effects on worker survival compared to the control.

**Table 2.**
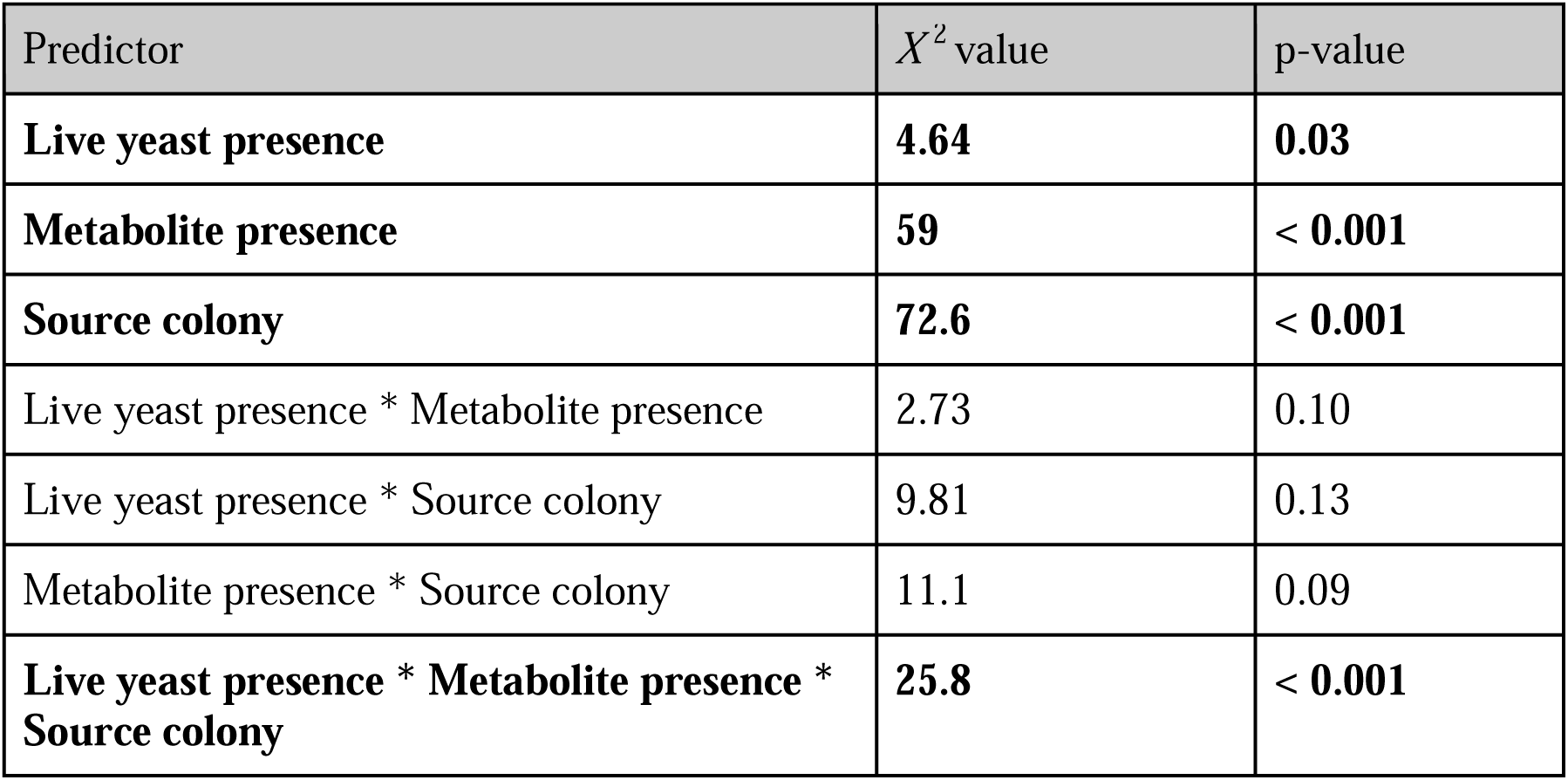
Effects of yeast treatments and source colony on survival of *B. impatiens* workers. Results are from a Cox proportional hazards model with microcolony ID as a random effect.

| Predictor | $X^2$ value | p-value |
| --- | --- | --- |
| **Live yeast presence** | **4.64** | **0.03** |
| **Metabolite presence** | **59** | **< 0.001** |
| **Source colony** | **72.6** | **< 0.001** |
| Live yeast presence * Metabolite presence | 2.73 | 0.10 |
| Live yeast presence * Source colony | 9.81 | 0.13 |
| Metabolite presence * Source colony | 11.1 | 0.09 |
| **Live yeast presence \* Metabolite presence \* Source colony** | **25.8** | **< 0.001** |

### Nectar consumption was unaffected by treatments, but differed across source colonies

Microcolonies consumed similar amounts of nectar regardless of live yeast presence (F_1,29_ = 2.21, p = 0.15) or metabolite presence (F_1,29_ = 0.001, p = 0.98). However, microcolonies from different source colonies consumed different amounts of nectar (F_4,29_ = 16.2, p < 0.001).

There were no significant interactions between any of the factors (p > 0.05 for all comparisons).

### Fungal treatments reduce Aspergillus prevalence, but may exacerbate negative pathogen effects

Because fungal pathogen infection was observed in a subset of microcolonies, colonies were screened for pathogens using PCR and qPCR. *Ascosphaera* was not detected in any microcolony, but *Aspergillus* was detected in 25 of 76 (33%) of microcolonies, and *A. flavus* was detected in 5 of 76 (6.6%) microcolonies using PCR. We further investigated *A. flavus* abundance using qPCR, and found that most microcolonies contained very low amounts, while a few microcolonies exhibiting symptomatic infection contained very high levels (range 0.09 - 3,533,021 copies, median = 8.5 copies).

Live yeast presence and metabolite presence interacted to impact *Aspergillus* presence, with control microcolonies exhibiting the greatest incidence (*X* ^2^ = 4.13, df = 1, p = 0.04, Figure 3A). Source colonies also differed in *Aspergillus* presence (*X* ^2^ = 24.6, df = 6, p < 0.001). However, the presence and abundance of known pathogen *A. flavus* (detected with much lower incidence) were not significantly affected by live yeast presence, metabolite presence, or their interaction (Table S4), though source colonies differed in both presence (*X* ^2^ = 17.3, df = 6, p = 0.008) and abundance (F_6,66_ = 8.11, p < 0.001) of *A. flavus*.

**Figure 3.**
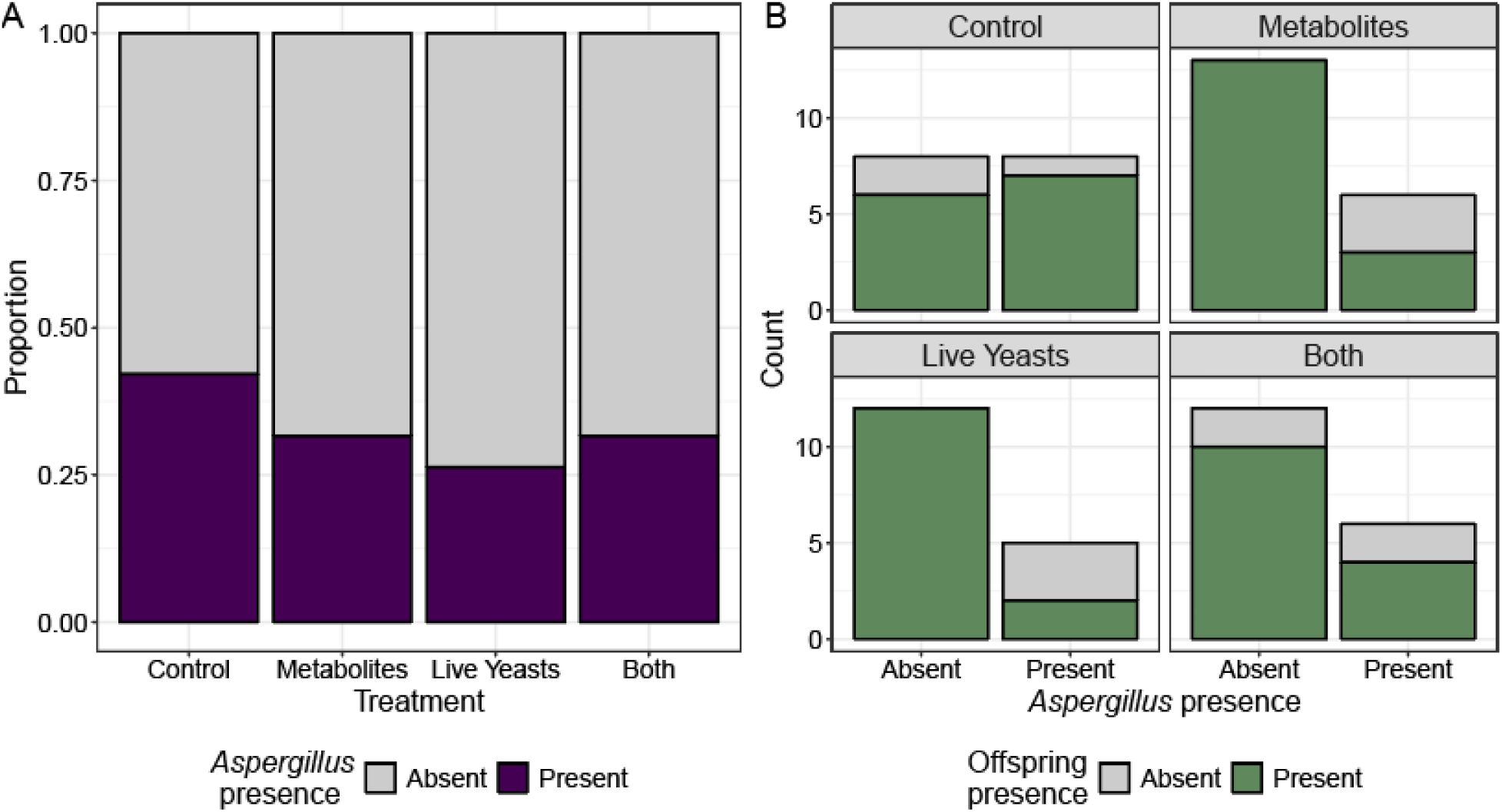
The prevalence of *Aspergillus* in microcolonies exposed to yeast treatments, and its impact on microcolony offspring production. A) Yeast treatments interacted to impact *Aspergillus* prevalence, with control microcolonies exhibiting the highest incidence (*X* ^2^ = 4.13, df = 1, p = 0.04). B) *Aspergillus* negatively impacted offspring production (*X* ^2^ = 7.71, df = 1, p = 0.006), though this effect also depended on yeast treatments (*X* ^2^ = 10.05, df = 1, p = 0.002). Panel A includes microcolonies from all source colonies, while Panel B includes microcolonies from source colonies that tested positive for *Aspergillus* (70 microcolonies).

Considering only source colonies that tested positive for *Aspergillus* (all source colonies except SS4), *Aspergillus* presence reduced the probability that a microcolony would produce offspring, although this effect was dependent on live yeast and metabolite presence and was most pronounced for microcolonies receiving live yeast cells or metabolites alone (*Aspergillus* presence * live yeast presence * metabolite presence interaction, *X* ^2^ = 10.05, df = 1, p = 0.002, Figure 3B). *Aspergillus* presence also interacted with live yeast presence and metabolite presence to impact total offspring number; microcolonies receiving no fungi or both live cells and metabolites did not respond as negatively to *Aspergillus* presence as those receiving either live cells or metabolites singly. However, this result was only marginally significant (three-way interaction, *X* ^2^ = 3.19, df = 1, p = 0.074). The presence of *Aspergillus* within a microcolony had no impact on adult bumble bee survival, and did not interact with any yeast treatments to impact survival (Logrank test, global p = 0.5).

### The presence of yeast cells alters chemical composition of bee diets

To determine how yeast treatments impacted the chemical composition of diets provided to bees, GC-MS metabolomics was performed on nectar and pollen taken from microcolonies three days after treatment application. Qualitatively, nectar contained a lower diversity of compounds than pollen (Figure S1) and contained mostly glucose, fructose, and sucrose present at high relative abundance. Pollen (here, a combination of nectar and pollen) contained mainly proline, glucose, fructose, gluconic acid lactone, and sorbitol. The diversity of compounds in nectar and pollen was unaffected by treatments (Table S5).

The presence of live yeast cells altered the chemical composition of nectar (RDA, F_1,16_ = 1.60, R^2^ = 0.06, p = 0.032, Figure 4A), while metabolite presence (F_1,16_ = 1.31, p = 0.10) and the interaction between treatments (F_1,16_ = 1.27, p = 0.12) had no effect. For pollen provisions, the interaction of live yeast presence and metabolite presence impacted compound composition (F_1,16_ = 1.51, R^2^ = 0.025, p = 0.042, Figure 4B).

**Figure 4.**
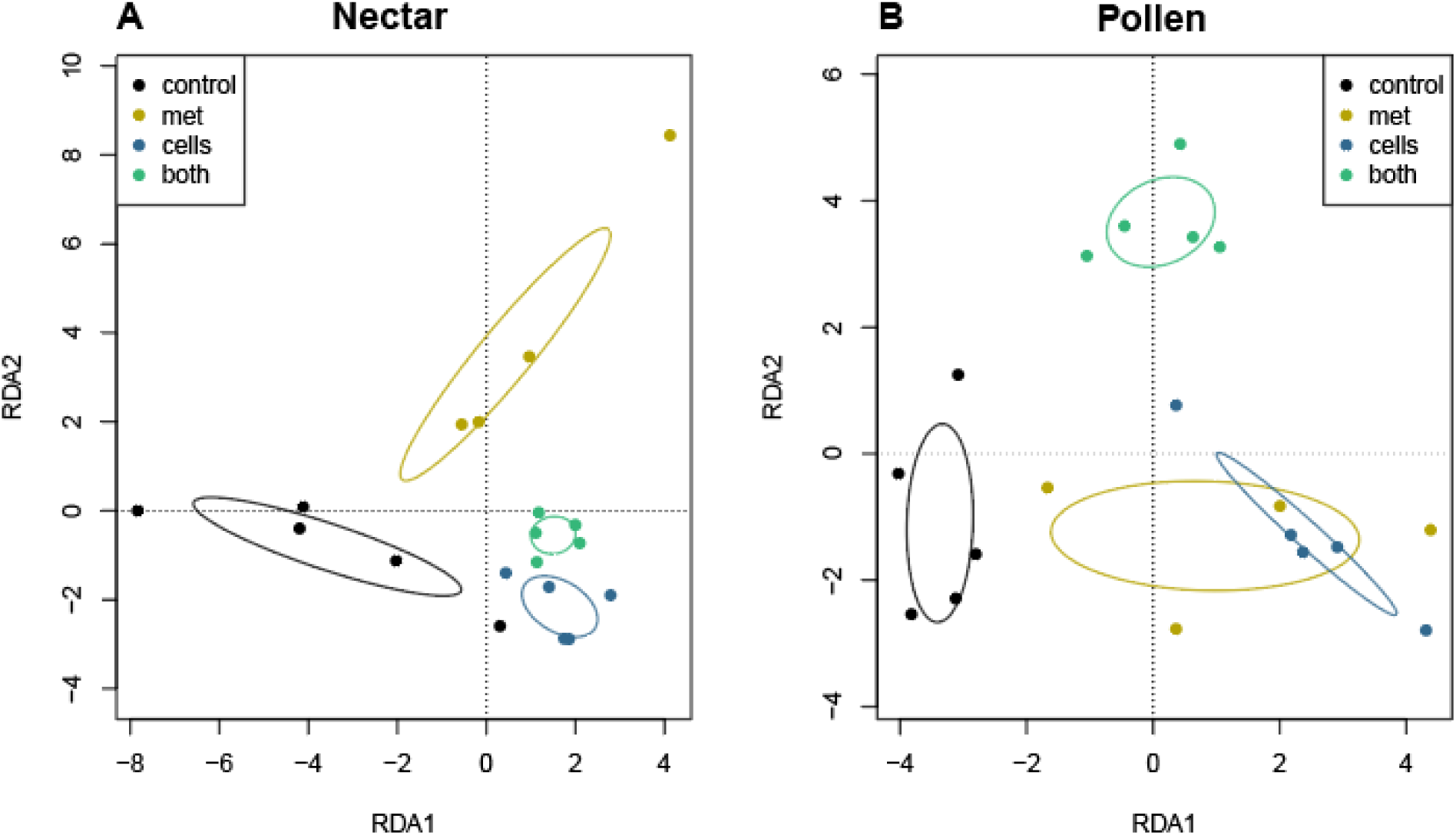
Ordination plots of metabolomics profiles of A) nectar and B) pollen provisions provided to *B. impatiens* microcolonies across different fungal treatments. Living yeast presence impacted the metabolomic composition of nectar samples (F_1,16_ = 1.60, R^2^ = 0.06, p = 0.032), while living yeast and metabolite presence interacted to impact pollen metabolites (F_1,16_ = 1.27, R^2^ = 0.025, p = 0.12).

Using live yeast presence as a response, random forest was able to successfully classify nectar samples to treatment based on chemical compound composition 90% of the time (10% OOB error). Using metabolite presence as a response, the classification accuracy was 75% (25% OOB error). For pollen samples, random forest correctly classified samples to live yeast treatments 60% of the time (40% OOB error) but only 25% of the time (75% OOB error) when metabolite presence was used as a predictor. Chemical features that were most important in distinguishing treatment groups differed by sample type, and were largely unidentified compounds (Figure S2).

In nectar, four compounds differed between metabolite presence treatments. Two unidentified compounds, 403585 (F_1,16_ = 103, p < 0.001) and 403592 (F_1,16_ = 120, p < 0.001) were lower in metabolite treatments, while unidentified compound 121482 was higher in metabolite treatments (F_1,16_ = 19.7, p = 0.03, Figure S3A). Adenosine was marginally lower in metabolite treatment groups (F_1,16_ = 15.5, p = 0.05). For pollen, only one unidentified compound, 112602, differed between treatments, and was lower in metabolite treatments (F_1,16_ = 57.3, p < 0.001, Figure S3B). Mass spectra of these unidentified compounds is provided in Table S6.

## Discussion

The experiment performed here tested several possible mechanisms through which symbiotic fungi might benefit bumble bees. Specifically, we tested if bees benefit from living fungal cells or from fungal metabolites and/or from alteration of food resources by fungi. We found no benefit of fungi across all microcolonies, either as living cells, their metabolites, or their effects on food, but we did observe treatment effects that were dependent on source colony identity, suggesting context-specific benefits of yeasts.

Colonies may have responded differently to fungal treatments for a variety of reasons, including differences in colony age, queen and worker health, genetic background, and variation in pre-existing microbial communities associated with each bee colony. Some studies document strain-level differences in bacterial gut microbiome composition between commercial *B. impatiens* source colonies (Hammer et al. 2022, Meeus et al. 2015; but see Rutkowski et al. 2022). In addition, microbial communities of stored provisions can vary between colonies, particularly in the presence and abundance of pathogens (Dharampal et al. 2020, Graystock et al. 2013), including viruses, *Vairimorpha, Crithidia*, and *Ascosphaera*, among others (Pereria et al. 2019, Strange et al. 2022). Previous work has documented the presence of *Ascosphaera* in the pollen provided to colonies (Dharampal et al. 2020, Rutkowski et al. 2022), which can infect adult bumble bees (Maxfield-Taylor et al. 2015). For this reason, we sterilized the pollen provided to microcolonies in the current experiment. Despite this, we observed offspring of microcolonies from one source colony (SS5) infected by the fungal pathogen *Aspergillus flavus*, and upon further investigation, observed variation in the presence of *Aspergillus* among source colonies. Although we did not manipulate the presence of this pathogen, variation in its presence allowed for a natural experiment testing yeast effects on bee-*Aspergillus* interactions. Interestingly, microcolonies derived from the source colony exhibiting symptomatic infection were the only ones that benefitted from a combination of live fungal cells and metabolites, though the presence and abundance of *A. flavus* was unaffected by fungal treatment (perhaps due to low replication within this source colony).

Results of other studies suggest that bee-associated fungi can suppress the growth of multiple bee pathogens and parasites: the presence of yeasts in *B. impatiens* colonies was associated with the absence of certain pathogenic fungi including *Ascosphaera apis* (Dharampal et al. 2020) and multiple yeasts suppress the growth of *Crithidia bombi* in vitro (Pozo et al. 2019). Our results add further experimental *in vivo* evidence of anti-pathogen effects of yeasts: here, any microcolonies treated with live yeasts or their metabolites had reduced incidence of *Aspergillus* compared to the control, suggesting possible suppression by symbiotic yeasts and their chemical byproducts. However, when *Aspergillus* was present in colonies, yeasts seemed to exacerbate negative effects of the pathogen, leading to lower rates of offspring production. Honeybees stressed with pesticides or pathogens more frequently hosted yeasts, suggesting that yeasts may exploit weakened bee hosts (Gilliam 1973, Gilliam et al. 1974). This may also be the case for bumble bees experiencing pressure from pathogens, but as this yeast-pathogen interaction was not the focus of this experiment, the conclusions we can draw from these data are limited, and further experiments explicitly testing the interactions of bee-associated yeasts with *Aspergillus* and other bee pathogens are necessary.

The current results add to a body of literature documenting variable and context-dependent impacts of fungi on bee health. Previous yeast addition experiments have found impacts of yeasts that vary based on bee and yeast species used. A previous study of *B. terrestris* used a similar study design to differentiate the effects of yeast cells and yeast metabolites/yeast-modified media on bee colonies (Pozo et al. 2020). They found that the nectar-associated yeast species *Metschnikowia gruessii* and *Rhodotorula mucilaginosa* as well as the bee-associated *Candida bombiphila* (=*Wickerhamiella bombiphila*) led to the production of more offspring, but this effect was independent of how the yeasts were administered. *M. gruessii, M. reukaufii*, and *C. bombiphila* also reduced larval mortality in these colonies. *Candida bombi* (=*Starmerella bombi*) was found to have no impact on colony health, as we found here for the related species *Starmerella sorbosivorans*. Another study that added the nectar yeast *Metschnikowia reukaufii* to *B. impatiens* microcolonies found it had no impact on colony development, though it was attractive to bees in foraging trials (Schaeffer et al. 2017). Finally, previous results using the same yeast community as in this study showed that fungal addition was beneficial to *B. impatiens* survival and reproduction (Rutkowski et al. 2022). In all, these results suggest that responses of bees to fungal supplementation are dependent on both bee and fungal identity, and that even if these are held constant, outcomes may still vary based on other underlying factors.

Given the known impacts of yeast metabolism on floral nectar traits, such as the concentration of amino acids and secondary metabolites, sugar concentration and composition, and volatile profile (Herrera et al. 2008, Rering et al. 2021, Vannette & Fukami 2016, Vannette & Fukami 2018), it is surprising that metabolomics analysis did not reveal large differences in compound composition between treatment groups for nectar or pollen. Most of the compounds that were detected as significantly different between treatments were unidentified, limiting the interpretation of our results. It is possible that yeasts alter metabolite composition in additional ways that were not detected using the untargeted primary metabolism method we used, which generated relative abundance measures of most compounds but not absolute abundances. Additionally, GC-MS mainly targets carbohydrates, amino acids, hydroxyl acids, free fatty acids, purines, pyrimidines, and aromatics. Other types of compounds, such as higher molecular weight lipids and complex carbohydrates, are not detected using this method and may be particularly important when considering yeast metabolism in this system. Two genera of yeasts used in this experiment, *Starmerella* and *Zygosaccharomyces*, produce lipids that have the potential to impact bee health. *Zygosaccharomyces* yeasts produce ergosterols necessary for development in certain stingless bee species (Paludo et al. 2018, Paula et al. 2023), while *Starmerella* yeasts produce sophorolipids that have broad antimicrobial properties (De Clercq et al. 2021, de O Caretta et al. 2022). Research investigating whether bee-associated yeasts produce these compounds in bee diets, and the resulting impacts on bee health, is necessary to further understand this bee-yeast association.

Overall, we found that impacts of yeast addition on *Bombus impatiens* microcolonies were inconsistent, and instead depended on source colony identity. These differences across source colonies may have been due to multiple factors, including variation in pathogen communities associated with each colony. We find partial support for the hypothesis that bee-associated yeasts may influence bee health through suppression of pathogens, as *Aspergillus* was more prevalent in control microcolonies. However, additional experiments explicitly testing this interaction are necessary to draw clear conclusions about the role of these yeasts in pathogen suppression and bee health. Taken together, we suggest that yeasts are common and perhaps ubiquitous symbionts of bumble bees and their stored food, but are likely not obligate mutualists, offering context-specific benefits as well as costs.

## Acknowledgements

We thank Jonathan Koch for sterilizing the pollen used in this experiment. We thank Richard Karban, Neal Williams, Shawn Christensen, Leta Landucci, Alexia Martin, and Dino Sbardellati for their comments which have improved this manuscript.

## Statements and Declarations

### Author Contributions

DR and RLV conceived of study. DR and MW carried out lab work and bee rearing. DR performed statistical analyses and wrote the first manuscript draft and DR and RLV contributed to revisions. All authors approved of the submission.

### Data availability statement

Bee response data and metabolomics data, along with all associated R code, is publicly available on Dryad (https://doi.org/10.5061/dryad.f1vhhmh5t)

## References

Bessette, E., & Williams, B. (2022). Protists in the Insect Rearing Industry: Benign Passengers or Potential Risk? Insects, 13(5), 482. 10.3390/insects13050482

Biedermann, P. H. W., & Vega, F. E. (2020). Ecology and Evolution of Insect–Fungus Mutualisms. Annual Review of Entomology, 65(1), 431–455. 10.1146/annurev-ento-011019-024910

Cerqueira, A. E. S., Hammer, T. J., Moran, N. A., Santana, W. C., Kasuya, M. C. M., & da Silva, C. C. (2021). Extinction of anciently associated gut bacterial symbionts in a clade of stingless bees. The ISME Journal 2021 15:9, 15(9), 2813–2816. 10.1038/s41396-021-01000-1

Christensen, S. M., Srinivas, S. N., Mcfrederick, Q. S., Danforth, B. N., Buchmann, S. L., Vannette, R. L., & Christensen, S. (2024). Symbiotic bacteria and fungi proliferate in diapause and may enhance overwintering survival in a solitary bee. The ISME Journal, 18(1), 89. 10.1093/ISMEJO/WRAE089

Cnaani, J., Schmid-Hempel, R., & Schmidt, J. O. (2002). Colony development, larval development and worker reproduction in Bombus impatiens Cresson. Insectes Sociaux, 49(2), 164–170. 10.1007/S00040-002-8297-8/METRICS

de Clercq, V., Roelants, S. L. K. W., Castelein, M. G., de Maeseneire, S. L., & Soetaert, W. K. (2021). Elucidation of the natural function of sophorolipids produced by starmerella bombicola. Journal of Fungi, 7(11), 917. 10.3390/JOF7110917/S1

de O Caretta, T., I Silveira, V. A., Andrade, G., Macedo, F., & P C Celligoi, M. A. (2022). Antimicrobial activity of sophorolipids produced by Starmerella bombicola against phytopathogens from cherry tomato. Journal of the Science of Food and Agriculture, 102(3), 1245–1254. 10.1002/JSFA.11462

de Paula, G. T., Melo, W. G. da P., Castro, I. de, Menezes, C., Paludo, C. R., Rosa, C. A., & Pupo, M. T. (2023). Further evidences of an emerging stingless bee-yeast symbiosis. Frontiers in Microbiology, 14, 1221724. 10.3389/FMICB.2023.1221724/BIBTEX

Dharampal, P. S., Carlson, C., Currie, C. R., & Steffan, S. A. (2019). Pollen-borne microbes shape bee fitness. Proceedings of the Royal Society B: Biological Sciences, 286(1904), 20182894. 10.1098/rspb.2018.2894

Dharampal, P. S., Diaz-Garcia, L., Haase, M. A. B., Zalapa, J., Currie, C. R., Hittinger, C. T., & Steffan, S. A. (2020). Microbial Diversity Associated with the Pollen Stores of Captive-Bred Bumble Bee Colonies. Insects, 11(4), 250. 10.3390/insects11040250

Echeverrigaray, S., Scariot, F. J., Foresti, L., Schwarz, L. V., Rocha, R. K. M., da Silva, G. P., Moreira, J. P., & Delamare, A. P. L. (2021). Yeast biodiversity in honey produced by stingless bees raised in the highlands of southern Brazil. International Journal of Food Microbiology, 347, 109200. 10.1016/j.ijfoodmicro.2021.109200

Evison, S. E., & Jensen, A. B. (2018). The biology and prevalence of fungal diseases in managed and wild bees. Current Opinion in Insect Science, 26, 105–113. 10.1016/J.COIS.2018.02.010

Feldhaar, H. (2011). Bacterial symbionts as mediators of ecologically important traits of insect hosts. Ecological Entomology, 36(5), 533–543. 10.1111/j.1365-2311.2011.01318.x

Garnier S., Ross N., Rudis R., Camargo A.P., Sciaini M., Scherer C. (2023). viridis(Lite) - Colorblind-Friendly Color Maps for R. viridis package version 0.6.3.

Gilliam, M. (1973). Are Yeasts Present in Adult Worker Honey Bees1 as a Consequence of Stress? Annals of the Entomological Society of America, 66(5), 1176–1176. 10.1093/aesa/66.5.1176

Gilliam, M., Wickerham, L. J., Morton, H. L., & Martin, R. D. (1974). Yeasts isolated from honey bees, Apis mellifera, fed 2,4-D and antibiotics. Journal of Invertebrate Pathology, 24, 349–356. 10.1016/0022-2011(74)90143-8

Gilliam, M. (1997). Identification and roles of non-pathogenic microflora associated with honey bees. FEMS Microbiology Letters, 155(1), 1–10. 10.1111/j.1574-6968.1997.tb12678.x

Graystock, P., Yates, K., Evison, S. E. F., Darvill, B., Goulson, D., & Hughes, W. O. H. (2013). The Trojan hives: pollinator pathogens, imported and distributed in bumblebee colonies. Journal of Applied Ecology, 50(5), 1207–1215. 10.1111/1365-2664.12134

Hammer, T. J., Easton-Calabria, A., & Moran, N. A. (2022). Microbiome assembly and maintenance across the lifespan of bumble bee workers. Molecular Ecology, 32(3), 724– 740. 10.1111/MEC.16769

Herrera, C. M., García, I. M., Pérez, R. (2008). Invisible floral larcenies: microbial communities degrade floral nectar of bumble bee-pollinated plants. Ecology, 89(9), 2369–2376. 10.1890/08-0241.1

Herrera, C. M., Pozo, M. I., & Medrano, M. (2013). Yeasts in nectar of an early-blooming herb: sought by bumble bees, detrimental to plant fecundity. Ecology, 94(2), 273–279. 10.1890/12-0595.1

Kaufman, M. G., Walker, E. D., Odelson, D. A., & Klug, M. J. (2000). Microbial Community Ecology & Insect Nutrition. American Entomologist, 46(3), 173–185. 10.1093/ae/46.3.173

Kwong, W. K., Medina, L. A., Koch, H., Sing, K. W., Soh, E. J. Y., Ascher, J. S., Jaffé, R., & Moran, N. A. (2017). Dynamic microbiome evolution in social bees. Science Advances, 3(3). 10.1126/sciadv.1600513

Leigh, B. A., Bordenstein, S. R., Brooks, A. W., Mikaelyan, A., & Bordenstein, S. R. (2018). Finer-Scale Phylosymbiosis: Insights from Insect Viromes. MSystems, 3(6). 10.1128/msystems.00131-18

Li, H., Young, S. E., Poulsen, M., & Currie, C. R. (2021). Symbiont-Mediated Digestion of Plant Biomass in Fungus-Farming Insects. Annual Review of Entomology, 66(Volume 66, 2021), 297–316. 10.1146/ANNUREV-ENTO-040920-061140/CITE/REFWORKS

Li, X., Wheeler, G. S., & Ding, J. (2012). A leaf-rolling weevil benefits from general saprophytic fungi in polysaccharide degradation. Arthropod-Plant Interactions, 6(3), 417–424. 10.1007/S11829-012-9194-3/FIGURES/3

Liaw A. & Wiener M. (2002). Classification and Regression by randomForest. R News 2(3), 18--22.

Lopez-Vaamonde, C., Brown, R. M., Lucas, E. R., Pereboom, J. J. M., Jordan, W. C., & Bourke, A. F. G. (2007). Effect of the queen on worker reproduction and new queen production in the bumble bee Bombus terrestris. Apidologie, 38(2), 171–180. 10.1051/APIDO:2006070

Maxfield-Taylor, S. A., Mujic, A. B., & Rao, S. (2015). First Detection of the Larval Chalkbrood Disease Pathogen Ascosphaera apis (Ascomycota: Eurotiomycetes: Ascosphaerales) in Adult Bumble Bees. PLOS ONE, 10(4), e0124868. 10.1371/journal.pone.0124868

Meeus, I., Parmentier, L., Billiet, A., Maebe, K., van Nieuwerburgh, F., Deforce, D., Wäckers, F., Vandamme, P., & Smagghe, G. (2015). 16S rRNA Amplicon Sequencing Demonstrates that Indoor-Reared Bumblebees (Bombus terrestris) Harbor a Core Subset of Bacteria Normally Associated with the Wild Host. PLOS ONE, 10(4), e0125152. 10.1371/JOURNAL.PONE.0125152

Mehdiabadi, N. J., & Schultz, R. (2009). Natural history and phylogeny of the fungus-farming ants (Hymenoptera: Formicidae: Myrmicinae: Attini). Myrmecological News, 13, 37–55.

Moran, N. A., Tran, P., & Gerardo, N. M. (2005). Symbiosis and insect diversification: An ancient symbiont of sap-feeding insects from the bacterial phylum Bacteroidetes. Applied and Environmental Microbiology, 71(12), 8802–8810. 10.1128/AEM.71.12.8802-8810.2005

Mueller, U. G., & Gerardo, N. (2002). Fungus-farming insects: Multiple origins and diverse evolutionary histories. In Proceedings of the National Academy of Sciences of the United States of America (Vol. 99, Issue 24, pp. 15247–15249). National Academy of Sciences. 10.1073/pnas.242594799

Nasir, H., & Noda, H. (2003). Yeast-like symbiotes as a sterol source in anobiid beetles (Coleoptera, Anobiidae): Possible metabolic pathways from fungal sterols to 7-dehydrocholesterol. Archives of Insect Biochemistry and Physiology, 52(4), 175–182. 10.1002/arch.10079

Noda, H., & Koizumi, Y. (2003). Sterol biosynthesis by symbiotes: Cytochrome P450 sterol C-22 desaturase genes from yeastlike symbiotes of rice planthoppers and anobiid beetles. Insect Biochemistry and Molecular Biology, 33(6), 649–658. 10.1016/S0965-1748(03)00056-0

Oksanen J, Simpson G, Blanchet F, Kindt R, Legendre P et al. (2022). vegan: Community Ecology Package. R package version 2.6–2, https://CRAN.R-project.org/package=vegan.

Paludo, C. R., Pishchany, G., Andrade-Dominguez, A., Silva-Junior, E. A., Menezes, C., Nascimento, F. S., Currie, C. R., Kolter, R., Clardy, J., & Pupo, M. T. (2019). Microbial community modulates growth of symbiotic fungus required for stingless bee metamorphosis. PLOS ONE, 14(7), e0219696. 10.1371/JOURNAL.PONE.0219696

Paludo, C. R., Menezes, C., Silva-Junior, E. A., Vollet-Neto, A., Andrade-Dominguez, A. et al. (2018). Stingless Bee Larvae Require Fungal Steroid to Pupate. Scientific Reports 2018 8:1, 8(1), 1–10. 10.1038/s41598-018-19583-9

Pang, Z., Chong, J., Zhou, G., de Lima Morais, D. A., Chang, L., Barrette, M., Gauthier, C., Jacques, P. É., Li, S., & Xia, J. (2021). MetaboAnalyst 5.0: Narrowing the gap between raw spectra and functional insights. Nucleic Acids Research, 49(W1), W388–W396. 10.1093/nar/gkab382

Pereira, K. de S., Meeus, I., & Smagghe, G. (2019). Honey bee-collected pollen is a potential source of Ascosphaera apis infection in managed bumble bees. Scientific Reports, 9(1). 10.1038/s41598-019-40804-2

Pozo, M. I., van Kemenade, G., van Oystaeyen, A., Aledón Catalá, T., Benavente, A., van den Ende, W., Wäckers, F., & Jacquemyn, H. (2020). The impact of yeast presence in nectar on bumble bee behavior and fitness. Ecological Monographs, 90(1). 10.1002/ecm.1393

Pozo, M. I., Mariën, T., van Kemenade, G., Wäckers, F., & Jacquemyn, H. (2021). Effects of pollen and nectar inoculation by yeasts, bacteria or both on bumblebee colony development. Oecologia, 195(3), 689–703. 10.1007/s00442-021-04872-4

R Core Team (2022). R: A language and environment for statistical computing. R Foundation for Statistical Computing, Vienna, Austria. URL https://www.R-project.org/.

Rering, C.C., Rudolph, A.B. and Beck, J.J. (2021). Pollen and yeast change nectar aroma and nutritional content alone and together, but honey bee foraging reflects only the avoidance of yeast. Environ Microbiol, 23: 4141–4150. 10.1111/1462-2920.15528

Rutkowski, D., Litsey, E., Maalouf, I., & Vannette, R. L. (2022). Bee-associated fungi mediate effects of fungicides on bumble bees. Ecological Entomology, 47(3), 411–422. 10.1111/een.13126

Rutkowski, D., Weston, M., & Vannette, R. L. (2023). Bees just wanna have fungi: a review of bee associations with nonpathogenic fungi. FEMS Microbiology Ecology, 99(8), 1–16. 10.1093/FEMSEC/FIAD077

Schaeffer, R. N., & Irwin, R. E. (2014). Yeasts in nectar enhance male fitness in a montane perennial herb. Ecology, 95(7), 1792–1798. 10.1890/13-1740.1

Schaeffer, R. N., Mei, Y. Z., Andicoechea, J., Manson, J. S., & Irwin, R. E. (2017). Consequences of a nectar yeast for pollinator preference and performance. Functional Ecology, 31(3), 613–621. 10.1111/1365-2435.12762

Scully, E. D., Geib, S. M., Hoover, K., Tien, M., Tringe, S. G., Barry, K. W., Glavina del Rio, T., Chovatia, M., Herr, J. R., & Carlson, J. E. (2013). Metagenomic Profiling Reveals Lignocellulose Degrading System in a Microbial Community Associated with a Wood-Feeding Beetle. PLoS ONE, 8(9), e73827. 10.1371/journal.pone.0073827

Smilde, A. K., Jansen, J. J., Hoefsloot, H. C. J., Lamers, R.-J. A. N., van der Greef, J., & Timmerman, M. E. (2005). ANOVA-simultaneous component analysis (ASCA): a new tool for analyzing designed metabolomics data. Bioinformatics, 21(13), 3043–3048. 10.1093/bioinformatics/bti476

Starmer, W. T., Barker, J. S. F., Phaff, H. J., & Fogleman, J. C. (1986). Adaptations of drosophila and yeasts: Their interactions with the volatile 2-propanol in the cactus–micro organism–drosophila model system. Australian Journal of Biological Sciences, 39(1), 69–77. 10.1071/BI9860069

Steffan, S. A., Dharampal, P. S., Danforth, B. N., Gaines-Day, H. R., Takizawa, Y., & Chikaraishi, Y. (2019). Omnivory in bees: Elevated trophic positions among all major bee families. The American Naturalist, 704281. 10.1086/704281

Steffan, S. A., Dharampal, P. S., Kueneman, J. G., Keller, A., Argueta-Guzmán, M. P. et al. (2023). Microbes, the ‘silent third partners’ of bee–angiosperm mutualisms. Trends in Ecology & Evolution, 0(0). 10.1016/J.TREE.2023.09.001

Strange, J. P., Colla, S. R., Duennes, M., Evans, E., Figueroa, L. L. et al. (2021). Developing a Commercial Bumble Bee Clean Stock Certification Program: A white paper of the North American Pollinator Protection Campaign Bombus Task Force. https://www.iucnredlist.org/

Strange, J. P., Tripodi, A. D., Huntzinger, C., Knoblett, J., Klinger, E. et al. (2023). Comparative analysis of 3 pollen sterilization methods for feeding bumble bees. Journal of Economic Entomology, 116(3), 662–673. 10.1093/JEE/TOAD036

Therneau T (2022). A Package for Survival Analysis in R. R package version 3.3–1, https://CRAN.R-project.org/package=survival.

Vanderpool, D., Bracewell, R. R., & McCutcheon, J. P. (2018). Know your farmer: Ancient origins and multiple independent domestications of ambrosia beetle fungal cultivars. Molecular Ecology, 27(8), 2077–2094. 10.1111/mec.14394

Vannette, R.L. & Fukami, T. (2016), Nectar microbes can reduce secondary metabolites in nectar and alter effects on nectar consumption by pollinators. Ecology, 97: 1410–1419. 10.1890/15-0858.1

Vannette, R.L. & Fukami, T. (2018) Contrasting effects of yeasts and bacteria on floral nectar traits, Annals of Botany, 121(7):1343–1349, 10.1093/aob/mcy032

Vega, F. E., & Dowd, P. F. (2005). The Role of Yeasts as Insect Endosymbionts. In Insect-Fungal Associations - Ecology and Evolution (pp. 211–243).

Velthuis, H. H. W., & van Doorn, A. (2006). A century of advances in bumblebee domestication and the economic and environmental aspects of its commercialization for pollination. In Apidologie (Vol. 37, Issue 4, pp. 421–451). EDP Sciences. 10.1051/apido:2006019

Venables, W. N. & Ripley, B. D. (2002) Modern Applied Statistics with S. Fourth Edition. Springer, New York. ISBN 0–387-95457-0

Wickham, H. (2016). ggplot2: Elegant Graphics for Data Analysis. Springer-Verlag New York.

Yang, M., Deng, G.-C., Gong, Y.-B., & Huang, S.-Q. (2019). Nectar yeasts enhance the interaction between Clematis akebioides and its bumblebee pollinator. Plant Biology, 21(4), 732–737. 10.1111/PLB.12957

Yun, J. H., Jung, M. J., Kim, P. S., & Bae, J. W. (2018). Social status shapes the bacterial and fungal gut communities of the honey bee. Scientific Reports, 8(1), 1–11. 10.1038/s41598-018-19860-7

