## Supplement for "Effects of live yeasts and their metabolic products on bumble bee microcolony development"

**Supplementary Material for**

**Effects of live yeasts and their metabolic products on bumble bee microcolony development**
Table S1. Sequencing results for fungal brood pathogens isolated from experimental microcolonies

| Source  microcolony | ITS2 sequence | Top BLAST hit | Percent  identity |
| --- | --- | --- | --- |
| M38 | TGAATCATCGAATCTTTGAACGCACATTGCGCCCCCTGGTATTCCGGGGGGCATGCCTGTCCGAGCGTCATTGCTGCCCATCAAGCACGGCTTGTGTGTTGGGTCGTCGTCCCCTCTCCGGGGGGGACGGGCCCCAAAGGCAGCGGCGGCACCGCGTCCGATCCTCGAGCGTATGGGGCTTTGTCACCCGCTCTGTAGGCCCGGCCGGCGCTTGCCGAACGCAAATCAATCTTTTTCCAGGTTGACCTCGGATCAGGTAGGGATACCMGCTGAACTTAAGCATAWCAAAAAGCGGA | *Aspergillus flavus* | 98.98 |
| M41 | TSAATCATCGAATCTTTGAACGCACATTGCGCCCCCTGGTATTCCGGGGGGCATGCCTGTCCGAGCGTCATTGCTGCCCATCAAGCACGGCTTGTGTGTTGGGTCGTCGTCCCCTCTCCGGGGGGGACGGGCCCCAAAGGCAGCGGCGGCACCGCGTCCGATCCTCGAGCGTATGGGGCTTTGTCACCCGCTCTGTAGGCCCGGCCGGCGCTTGCCGAACGCAAATCAATCTTTTTCCAGGTTGACCTCGGATCAGGTAGGGATACCCGCTGAACTTAAGCATAWCAAAAAGCGGAGGAA | *Aspergillus flavus* | 98.99 |

Table S2. Details of custom plasmid used to calculate *Aspergillus* *flavus* gene copy number from qPCR data

| Vector | Inserted sequence |
| --- | --- |
| Blue Heron pUC | CCCCCTGGTATTCCGGGGGGCATGCCTGTCCGAGCGTCATTGCTGCCCATCAAGCACGGCTTGTGTGTTGGGTCGTCGTCCCCTCTCCGGGGGGGACGGGCCCCAAAGGCAGCGGCGGCACCGCGTCCGATCCTCGAGCGTATGGGGCTTTGTCACCCGCTCTGTAGGCCCGGCCGGCGCTTGCCGAACGCAAATCAATCTTTTTCCAGGTTGACCTCGGATCAGGTAGGGATACCCGCTGAACTTAAGCAT |

Table S3. Effects of yeast treatments and source colony on microcolony offspring mass, separated by life stage.

| Response variable | Predictor variable | F-value | p-value |
| --- | --- | --- | --- |
| Egg mass | Live yeast presence | 0.018 | 0.90 |
|  | Metabolite presence | 1.09 | 0.31 |
|  | Source colony | 1.69 | 0.19 |
|  | Live yeast presence * Metabolite presence | 0.90 | 0.36 |
|  | Live yeast presence * Source colony | 0.58 | 0.57 |
|  | Metabolite presence * Source colony | 0.75 | 0.49 |
|  | Live yeast presence * Metabolite presence * Source colony | 1.38 | 0.28 |
| Larvae mass | Live yeast presence | 0.004 | 0.95 |
|  | **Metabolite presence** | **5.80** | **0.02** |
|  | Source colony | 0.75 | 0.61 |
|  | Live yeast presence * Metabolite presence | 0.010 | 0.92 |
|  | Live yeast presence * Source colony | 0.35 | 0.88 |
|  | Metabolite presence * Source colony | 0.61 | 0.72 |
|  | Live yeast presence * Metabolite presence * Source colony | 0.25 | 0.91 |
| Pupae mass | Live yeast presence | 0.21 | 0.68 |
|  | Metabolite presence | 0.00 | 0.98 |
|  | Source colony | 2.78 | 0.21 |
|  | Live yeast presence * Metabolite presence | 3.93 | 0.14 |
|  | Live yeast presence * Source colony | 0.92 | 0.79 |
|  | Metabolite presence * Source colony | 0.098 | 0.78 |
| Male mass | Live yeast presence | 2.07 | 0.18 |
|  | Metabolite presence | 0.17 | 0.69 |
|  | Source colony | 2.29 | 0.10 |
|  | Live yeast presence * Metabolite presence | 2.17 | 0.17 |
|  | Live yeast presence * Source colony | 2.43 | 0.12 |
|  | Metabolite presence * Source colony | 3.97 | 0.07 |

Table S4. Results of analyses of *A. flavus* presence and abundance across treatments and source colonies

| Response Variable | Predictor | Test | Test statistic | p-value |
| --- | --- | --- | --- | --- |
| *A. flavus* presence | Live yeast presence | Chi square | 0.30 | 0.59 |
|  | Metabolite presence | Chi square | 0.30 | 0.59 |
|  | Live yeast presence * Metabolite presence | Chi square | 0.24 | 0.62 |
|  | **Source colony** | **Chi square** | **17.3** | **0.008** |
| *A. flavus* abundance | Live yeast presence | F test | 0.0047 | 0.95 |
|  | Metabolite presence | F test | 1.43 | 0.24 |
|  | Live yeast presence * metabolite presence | F test | 0.0002 | 0.99 |
|  | **Source colony** | **F test** | **8.11** | **< 0.001** |

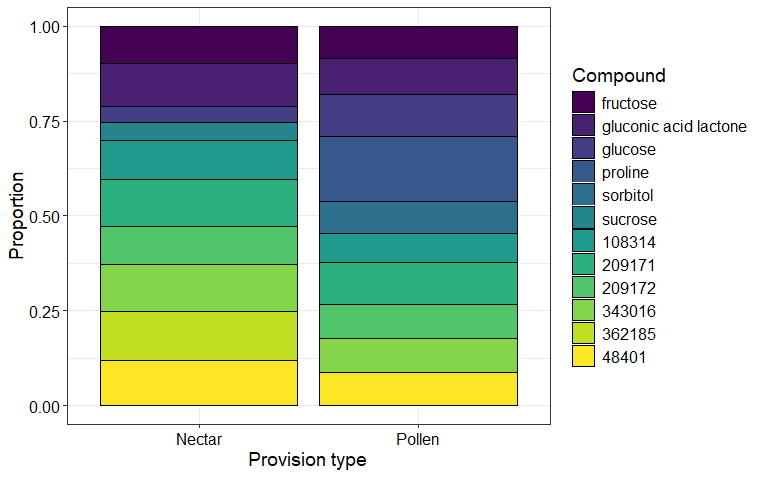

Figure S1. Top 10 compounds detected in nectar and pollen provisions using untargeted metabolomics with GC-TOF. Both pollen and nectar were dominated by sugars and sugar derivatives. Many top compounds were unidentified by the BinBase database, and their BinBase identifiers are reported here. Values represent non-normalized metabolomic data.

Table S5. Effects of treatments on nectar and pollen chemical compound richness

| Response Variable | Predictor | F-value | p-value |
| --- | --- | --- | --- |
| Nectar compound richness | Live yeast presence | 1.49 | 0.24 |
|  | Metabolite presence | 0.42 | 0.53 |
|  | Live yeast presence * Metabolite presence | 1.81 | 0.20 |
| Pollen compound richness | Live yeast presence | 0.22 | 0.65 |
|  | Metabolite presence | 0.037 | 0.85 |
|  | Live yeast presence * Metabolite presence | 2.46 | 0.14 |

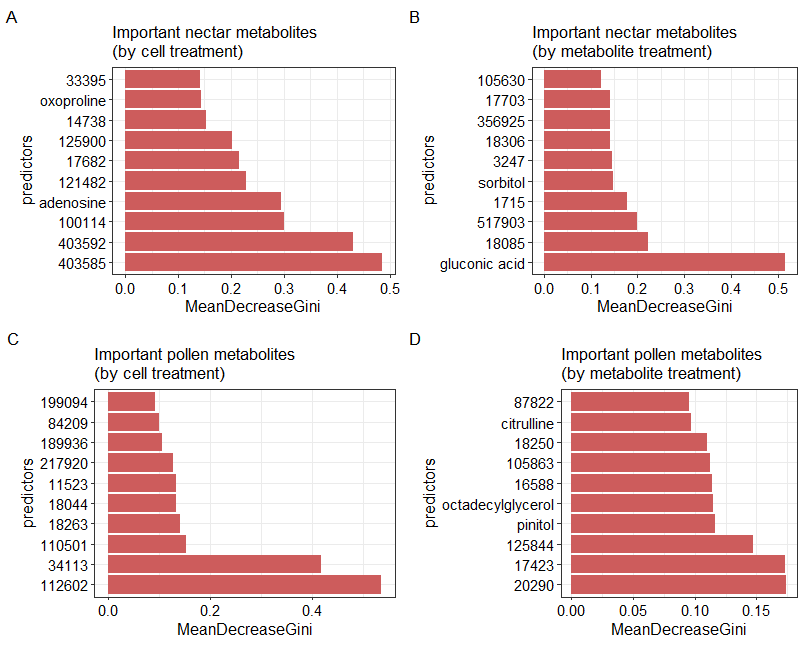

Figure S2. Chemical compounds that were important predictors of treatments in random forest analysis for nectar samples (A,B) and pollen samples (C,D). Many compounds were unidentified by the BinBase database, the BinBase identifiers of these compounds are reported here.

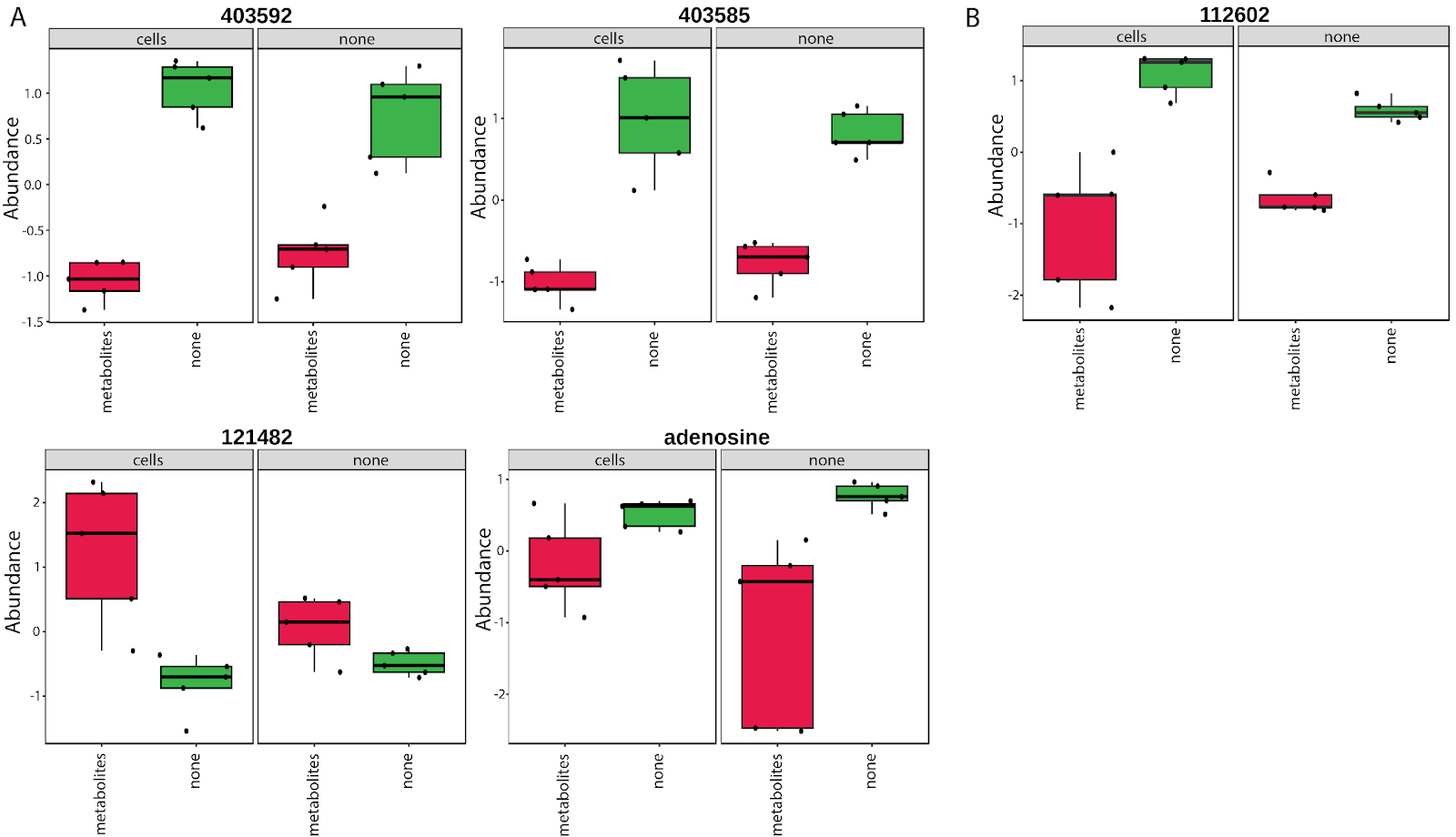

Figure S3. Chemical compounds that differed between yeast treatment groups in A) nectar and B) pollen according to ANOVA. For nectar, adenosine (F = 15.5, p = 0.05), as well as unidentified compounds 403592 (F = 120, p < 0.001) and 403585 (F = 103, p < 0.001) were lower in metabolite treatments, while unidentified compound 121482 (F = 19.7, p = 0.03) was higher in metabolite treatments. For pollen, unidentified compound 112602 (F = 57.3, p < 0.001) was lower in metabolite treatments. For compounds that were unidentified by the BinBase database, the BinBase identifiers of these compounds are reported here.

Table S6. Mass spectra of unidentified compounds that differed between treatment groups for pollen and nectar provisions

| Compound ID (BinBase) | Mass spectra (size:peak intensity) |
| --- | --- |
| 403585 | 89:2203.0 90:161.0 94:187.0 95:649.0 96:344.0 98:255.0 100:149.0 101:751.0 103:7990.0 104:790.0 105:540.0 108:49.0 109:29.0 111:352.0 112:53.0 122:39.0 124:295.0 125:9938.0 126:736.0 127:127.0 133:477.0 134:111.0 138:341.0 139:617.0 140:325.0 142:97.0 144:84.0 147:1128.0 148:237.0 152:325.0 154:556.0 155:491.0 167:50.0 168:1430.0 169:144.0 170:80.0 180:106.0 181:78.0 182:172.0 183:357.0 191:418.0 192:93.0 196:300.0 198:327.0 199:106.0 212:122.0 213:211.0 214:666.0 215:96.0 228:134.0 229:105.0 241:28.0 245:75.0 253:21.0 256:620.0 257:76.0 258:85.0 259:20.0 270:38.0 285:41.0 286:123.0 317:141.0 320:21.0 |
| 403592 | 85:710.0 86:91.0 87:96.0 88:305.0 89:4207.0 90:307.0 91:699.0 92:177.0 93:160.0 94:232.0 95:195.0 96:506.0 97:619.0 98:498.0 100:294.0 101:1151.0 103:12718.0 104:1267.0 105:661.0 106:109.0 108:69.0 110:688.0 111:682.0 112:363.0 113:268.0 114:58.0 115:831.0 119:363.0 124:262.0 125:8761.0 126:1252.0 127:61.0 128:53.0 133:3513.0 134:257.0 135:376.0 136:49.0 138:429.0 139:645.0 140:618.0 142:127.0 144:135.0 147:2239.0 148:211.0 152:573.0 153:39.0 154:491.0 155:640.0 156:511.0 159:376.0 163:74.0 164:73.0 168:906.0 170:197.0 175:142.0 177:130.0 180:89.0 182:104.0 183:330.0 186:501.0 191:364.0 193:295.0 194:90.0 196:193.0 198:467.0 199:107.0 200:103.0 202:68.0 205:662.0 206:170.0 212:95.0 214:463.0 219:124.0 220:155.0 221:107.0 228:230.0 229:79.0 230:141.0 232:69.0 233:48.0 240:63.0 242:31.0 243:667.0 244:206.0 256:485.0 257:270.0 258:145.0 259:45.0 267:622.0 268:210.0 274:59.0 282:185.0 286:133.0 302:92.0 317:130.0 318:29.0 328:51.0 332:14.0 335:38.0 343:22.0 348:30.0 354:15.0 365:18.0 376:10.0 379:24.0 391:10.0 392:26.0 406:21.0 407:15.0 414:11.0 434:7.0 436:16.0 446:7.0 447:19.0 449:18.0 452:23.0 454:12.0 455:27.0 465:21.0 473:13.0 474:18.0 475:22.0 477:26.0 480:21.0 493:15.0 500:19.0 |
| 121482 | 85:212.0 87:150.0 89:2092.0 90:382.0 95:250.0 99:410.0 100:756.0 101:2873.0 102:1004.0 103:9181.0 104:832.0 105:1698.0 106:175.0 108:51.0 112:67.0 113:296.0 114:410.0 115:827.0 116:1545.0 117:3454.0 118:342.0 119:241.0 124:36.0 126:27.0 127:66.0 128:171.0 129:5613.0 130:1315.0 131:2250.0 132:351.0 133:3979.0 134:213.0 135:346.0 140:129.0 141:696.0 142:519.0 143:1577.0 144:250.0 145:375.0 147:15323.0 148:2181.0 149:2073.0 150:298.0 151:107.0 152:32.0 153:166.0 155:467.0 156:47.0 157:1569.0 158:367.0 159:256.0 160:5695.0 161:1010.0 162:312.0 163:359.0 164:60.0 167:37.0 169:1631.0 170:243.0 171:192.0 172:80.0 173:203.0 175:217.0 177:195.0 178:112.0 180:69.0 181:88.0 182:10.0 183:99.0 184:70.0 185:136.0 186:269.0 189:1874.0 190:542.0 191:3480.0 192:591.0 197:76.0 199:375.0 200:109.0 201:206.0 202:45.0 203:477.0 204:12680.0 205:4865.0 206:1602.0 209:110.0 210:94.0 215:485.0 216:477.0 217:9407.0 218:2417.0 219:1230.0 220:217.0 221:272.0 223:110.0 227:152.0 228:421.0 229:443.0 230:397.0 231:927.0 232:263.0 233:293.0 234:119.0 240:63.0 241:55.0 242:82.0 243:1793.0 244:636.0 245:444.0 246:363.0 247:367.0 248:170.0 254:130.0 255:134.0 256:22.0 257:593.0 258:115.0 259:632.0 260:231.0 262:127.0 263:602.0 264:131.0 268:112.0 269:262.0 270:177.0 271:1066.0 272:288.0 273:300.0 274:119.0 275:66.0 276:224.0 277:267.0 279:27.0 290:49.0 291:458.0 292:495.0 293:155.0 294:79.0 297:24.0 299:17.0 300:853.0 301:179.0 302:112.0 303:66.0 304:41.0 305:580.0 306:257.0 307:890.0 308:358.0 309:165.0 311:11.0 315:12.0 319:722.0 320:141.0 321:236.0 322:89.0 331:266.0 332:216.0 333:114.0 334:20.0 335:27.0 343:63.0 345:7.0 346:54.0 347:130.0 348:51.0 349:48.0 350:41.0 353:17.0 358:70.0 360:415.0 361:4694.0 362:2309.0 363:888.0 364:422.0 365:85.0 366:51.0 388:71.0 389:85.0 390:497.0 391:303.0 392:112.0 393:78.0 400:144.0 417:40.0 418:25.0 434:28.0 435:67.0 436:567.0 437:2441.0 438:1635.0 439:694.0 440:297.0 441:34.0 450:72.0 451:186.0 452:176.0 453:50.0 475:80.0 479:92.0 480:338.0 481:242.0 482:133.0 483:30.0 490:30.0 491:100.0 |
| 112602 | 85:182.0 87:121.0 89:211.0 99:100.0 101:221.0 103:2762.0 104:110.0 105:22.0 106:9.0 111:174.0 117:397.0 127:205.0 129:1810.0 131:242.0 133:716.0 135:135.0 142:79.0 143:735.0 144:32.0 147:2115.0 148:331.0 152:47.0 154:72.0 155:122.0 168:23.0 171:32.0 177:105.0 190:29.0 191:61.0 195:36.0 201:22.0 202:24.0 215:18.0 217:9445.0 218:1714.0 219:648.0 220:38.0 225:26.0 242:45.0 243:172.0 244:75.0 245:73.0 247:110.0 258:72.0 260:24.0 314:28.0 319:23.0 332:46.0 335:25.0 336:18.0 349:17.0 351:46.0 357:25.0 364:22.0 369:46.0 405:34.0 408:21.0 410:27.0 421:27.0 442:16.0 447:24.0 454:18.0 461:18.0 472:17.0 476:18.0 489:17.0 491:19.0 |
